# Tumor irradiation combined with vascular-targeted photodynamic therapy enhances anti-tumor effects in preclinical prostate cancer

**DOI:** 10.1101/2020.11.26.400283

**Authors:** Hanna T Sjoberg, Yiannis Philippou, Anette L Magnussen, Iain DC Tullis, Esther Bridges, Andrea Chatrian, Joel Loefebvre, Ka Ho Tam, Emma A Murphy, Jens Rittscher, Dina Preise, Lilach Agemy, Tamar Yechezkel, Sean C Smart, Paul Kinchesh, Stuart Gilchrist, Danny P Allen, David A Scheiblin, Stephen J Lockett, David A Wink, Alastair D Lamb, Ian G Mills, Adrian Harris, Ruth J Muschel, Boris Vojnovic, Avigdor Scherz, Freddie C Hamdy, Richard J Bryant

## Abstract

**Rationale:** There is an important clinical need to improve the treatment of high risk localized and locally advanced prostate cancer (PCa), and to reduce the side effects of these treatments. We hypothesized that multi-modality therapy combining radiotherapy and vascular-targeted photodynamic therapy (VTP) could PCa tumour control compared against monotherapy with each of these treatments alone. This could provide proof-of-concept to take to the clinic. VTP is a focal therapy for localized PCa, which rapidly destroys targeted tumors through vascular disruption. Tumor vasculature is characterized by vessel immaturity, increased permeability, aberrant branching and inefficient flow. Fractionated radiotherapy (FRT) alters the tumor microenvironment and promotes transient vascular normalization.

**Objective:** We investigated whether sequential delivery of FRT followed by VTP 7 days later improves PCa tumor control compared to monotherapy with FRT or VTP alone.

**Findings:** FRT induced vascular normalization changes in PCa flank tumor allografts, improving vascular function as demonstrated using dynamic contrast enhanced magnetic resonance imaging. FRT followed by VTP significantly delayed tumor growth in flank PCa allograft pre-clinical models, compared with monotherapy with FRT or VTP alone, and improved overall survival.

**Conclusion:** Taken together, these results suggest that combining FRT and VTP could become a promising multimodal clinical strategy in PCa therapy. This provides proof-of-concept for this multi-modality therapy approach to take forward to early phase clinical trials.

## Introduction

There is an important unmet clinical need to improve the treatment outcomes for high risk prostate cancer (PCa) (which includes high-grade localized and locally advanced disease) (1–4), and to reduce the side effects of these treatments. Fractionated radiotherapy (FRT) combined with androgen deprivation therapy (ADT) is a curative option for high-risk PCa, however a third of cases recur posttreatment (1,4–10), and subsequent curative options are limited. Moreover, FRT and ADT have significant short- and long-term side effects (4,6). Whilst radical prostatectomy is a treatment option for high-risk PCa (11,12,21,13–20), there are unmet clinical needs to increase cure rates, and to reduce treatment toxicity.

Vascular-targeted photodynamic therapy (VTP) is a novel minimally invasive focal ablation surgical procedure, achieved by rapid free radical-mediated destruction of the tumor vasculature leading to focal necrosis (22–25). VTP is effective in the focal ablation of low-risk PCa (26–28), and has been investigated as salvage therapy for patients with radio-recurrent PCa (29,30), however to date it has not been investigated in combination with any other treatment as a multi-modality therapy approach to high-risk PCa. Moreover, it has not been investigated as a treatment option for high-risk PCa. VTP involves intravenous administration of the soluble photosensitizing agent WST-11, which conjugates to albumin and is focally activated with near-infrared illumination in the presence of oxygen. VTP requires the presence of functional tumor vasculature in order for the WST-11 photosensitizer to be efficiently delivered to the target tumor tissue prior to activation by locally applied near-infrared illumination and subsequent vascular and tumor destruction.

The tumor vasculature is typically characterized by blood vessel immaturity, increased permeability, increased interstitial pressure, absence of peri-vascular supporting cells, aberrant vessel branching and inefficient blood flow. These properties can impair therapeutic drug delivery and reduce the efficacy of treatment (31,32). FRT can alter several aspects of the tumor microenvironment, including the transient restoration of tumor blood vessel function, termed vascular normalization (33,34). Vascular normalization is characterized by inhibition of angiogenesis, restoration of peri-vascular cell support of tumor vessels, reduced vessel branching, and restoration of blood flow (35). These transient vascular changes may influence the effectiveness of VTP, and whilst VTP has been investigated in small observation cohort studies of patients with radio-recurrent PCa, to date the application of VTP during the period of transient vascular normalization post-FRT has not been investigated.

We aimed to investigate the combination of FRT and VTP as a multi-modality therapy approach to treating high-risk PCa in a pre-clinical model, in order to provide proof-of-concept of this approach to potentially inform early phase clinical trials. We specifically tested the hypothesis that sequential delivery of FRT followed 7 days later by VTP, with use of VTP during a period of transient post-FRT vascular normalization, may improve tumor control compared to monotherapy with FRT or VTP alone.

## Results

### VTP monotherapy induces delayed tumor growth in prostate cancer flank tumor allografts

We have reported previously the anti-tumor effects of FRT in a murine syngeneic immunocompetent TRAMP-C1 flank PCa tumor allograft model (36). In order to similarly assess the anti-tumor effects of VTP in this model, and subsequently combine FRT and VTP as multimodality therapy, we developed an enclosed optical irradiation system to deliver VTP to flank allograft tumors (**Figure 1**). A bespoke cradle was constructed to accommodate the anaesthetized animal in this optical system. Using this VTP platform, flank TRAMP-C1 tumor allografts were treated when they reached 100 mm^3^ volume with 9 mg/kg WST-11 at 120 mW/cm^2^ for 600 seconds (**Figure 2**). Treatment of TRAMP-C1 tumor allografts with 9 mg/kg resulted in tumor growth delay to a final tumor size of 400 mm^3^ compared with untreated control tumors (**Figures 2A-D**), with no significant welfare implications (**Figure 2E**), however all tumors eventually recurred.

**Figure 1.**
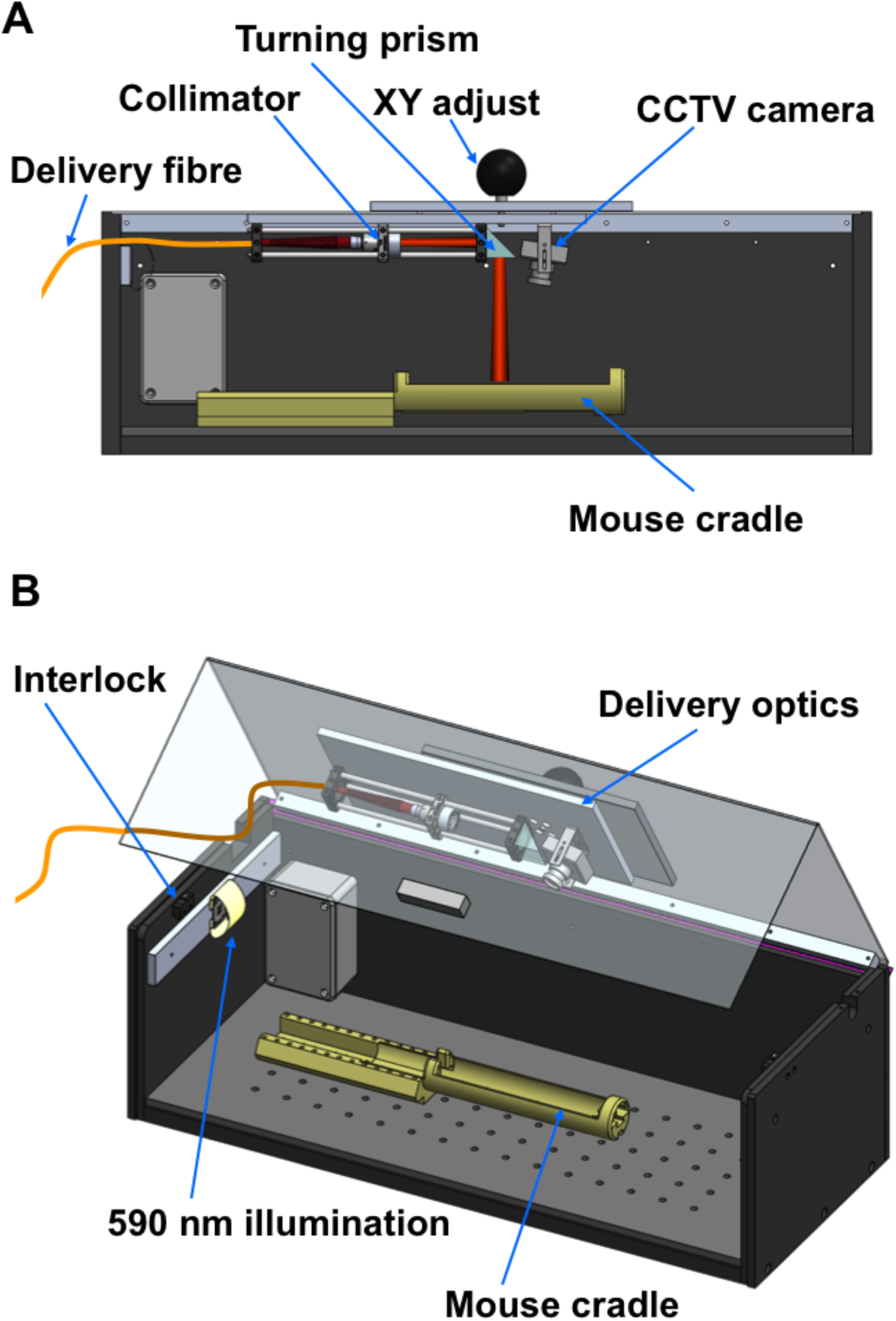
SolidWorks^®^ models used in the design of the VTP optical delivery enclosure. (**A**) View from the front with the front plate transparent to allow internal viewing of the enclosure contents. Homogenized excitation light is delivered via a multimode fibre coupled to a collimator, which delivers a slightly diverging light beam, with spot size adjustment achieved by changing the distance between collimator and prism. A turning prism directs light onto the flank tumor surface. A high dynamic range camera allows viewing of the beam and mouse, which is placed in a cradle. Details of the anesthesia system are not shown for clarity. (**B**) View of the enclosure with the lid slightly open to reveal the animal illumination, laser interlock switch and details of the adjustable cradle. The whole optical delivery system can be moved laterally to ensure correct illumination of the flank tumor.

**Figure 2.**
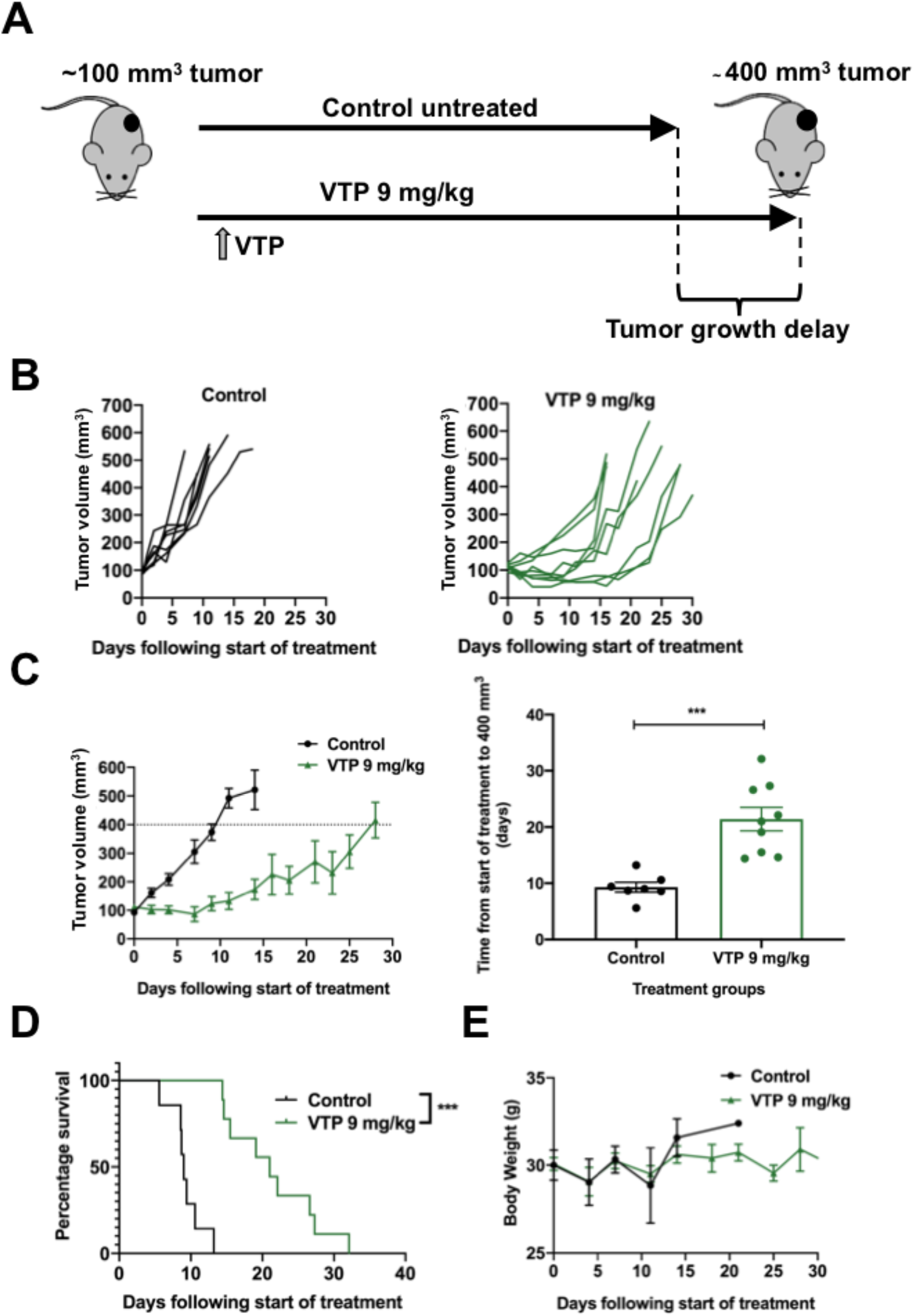
VTP causes tumor growth delay in TRAMP-C1 flank tumor allografts. (**A**) Outline schematic of treatment of subcutaneous flank TRAMP-C1 tumor allografts with VTP. (**B**) Growth kinetics of TRAMP-C1 tumors following indicated treatments (n = 7 untreated control; n = 10 VTP 9 mg/kg WST-11, 120 mW/cm^2^, 600 seconds). (**C**) Tumor growth delay to ≥400 mm^3^ analysis of TRAMP-C1 allograft tumors treated with VTP 9 mg/kg WST-11, 120 mW/cm^2^, 600 seconds. Data are presented as mean tumor volume ± SEM and analysed using ordinary one-way ANOVA with Tukey’s *post hoc* adjustment for multiple comparisons. \**p* < 0.05; \*\**p* < 0.01; *** *p* < 0.001, **** *p* < 0.0001. (**D**) Treatment of TRAMP-C1 tumor allografts with 9 mg/kg WST-11, 120 mW/cm^2^, for 600 seconds resulted in enhanced survival to a tumor size of 400 mm^3^ compared with untreated control tumors. (**E**) Median (range) body weight at start of experiment: untreated control = 21.2 g (20.2–23.5 g); VTP 9 mg/kg = 20.9 g (18.5–23.8 g).

### Radiotherapy induces vascular normalization in prostate cancer flank tumor allografts

To assess the potential for FRT to induce vascular changes in PCa tumors *in vivo* that might influence VTP treatment, flank TRAMP-C1 tumor allografts were treated with 3 x 5Gy FRT at 100 mm^3^ and harvested at 7 days (early) or at 400 mm^3^ tumor regrowth end-point (late) following initiation of treatment (**Figure 3A**). CD31 and αSMA expression was analysed in tissue sections from control untreated and FRT treated TRAMP-C1 tumors (**Figure 3B**). Image segmentation analysis revealed a reduced proportion of smaller-diameter CD31-positive vessels at 7 days post-initiation of FRT (early time-point) versus untreated control tumors (**Figure 3C**), whilst no difference was observed at ≥400 mm^3^ tumor recurrence post-FRT (late time point). The density of CD31-positive tumor vessels (or number of vessels/cm^2^) was lower for FRT-treated tumors than for untreated control tumors at 7 days (early time point) (**Figure 3D**). Two-dimensional Euclidian distance transformation analysis of the distance between any tissue image pixel and the closest CD31-positive vessel segment revealed a reduced frequency of short distances between any pixel and the closest CD31-positive vessel at 7 days post-initiation of FRT versus untreated control tumors (**Figure 3E**). This demonstrates that there were longer distances between CD31-positive vessels at 7 days post-initiation of FRT compared to control untreated tumors. No difference was observed at ≥400 mm^3^ tumor recurrence post-FRT.

**Figure 3.**
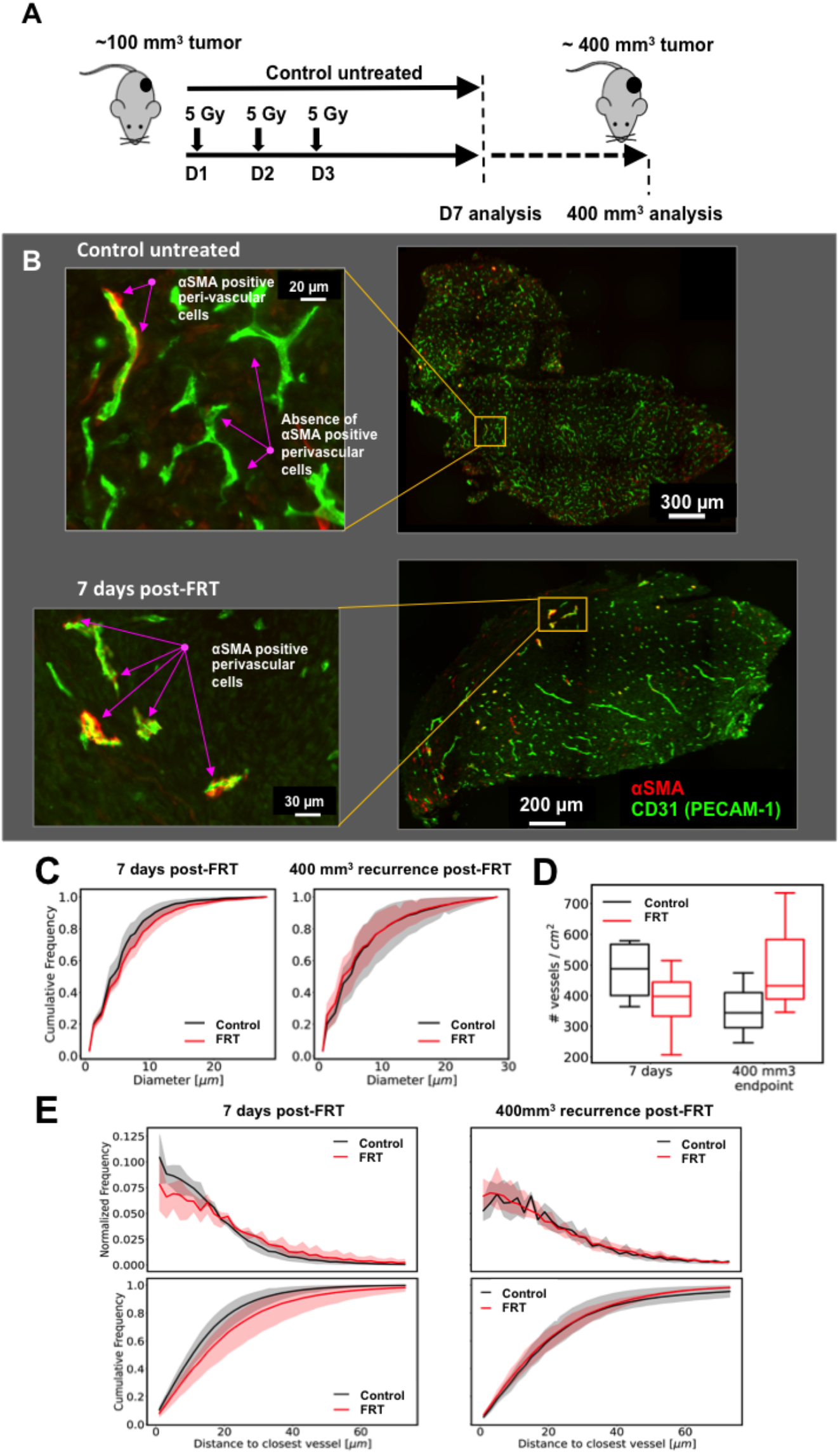
FRT induces vascular normalization in flank TRAMP-C1 PCa tumor allografts. (**A**) Outline schematic of treatment of tumors with 3 x 5 Gy FRT ahead of histological analysis of vascular changes. (**B-C**) Image segmentation analysis of immunofluorescence images from untreated control (n=3) and FRT-treated (n=5) TRAMP-C1 flank tumors (green – anti-CD31; red – anti-αSMA) revealed a reduced proportion of smaller-diameter blood vessels at 7 days postinitiation of FRT (early time point) compared to control; no difference was observed at the tumor regrowth to ≥400 mm^3^ endpoint (late time point). (**D**) The density (number of vessels/cm^2^) of CD31-positive tumor vessels was lower for FRT-treated tumors (n=5) than for control tumors (n=3) at 7 days (early time point). (**E**) 2D Euclidian distance transformation analysis of the distance between any tissue section pixel and the closest CD31-positive tumor vessel segment revealed differences between FRT-treated and control TRAMP-C1 tumors. This analysis revealed a reduced frequency of short distances between any tissue pixel and the closest CD31-positive tumor vessel at 7 days post-initiation of FRT (early time point), indicating that there were longer distances between tumor vessels at this time point. No difference was observed between FRT-treated and control TRAMP-C1 tumors at regrowth to ≥400 mm^3^ (late time point).

The association of digitally annotated αSMA-positive peri-vascular cells with CD31-positive vessels was then analyzed in TRAMP-C1 flank tumor allografts following FRT (**Figure 4A**). An increased fraction of CD31-positive tumor vessels with ≥1 adjacent αSMA-positive peri-vascular cell was observed in TRAMP-C1 flank tumor allografts at 7 days post-initiation of FRT compared to untreated control tumors (**Figure 4B**). This effect was not seen at the ≥400 mm^3^ tumor regrowth end-point post-FRT. An increased mean fraction of CD31-positive vessels with an adjacent αSMA-positive perivascular cell was observed in TRAMP-C1 flank tumor allografts at 7 days post-initiation of FRT compared to untreated control tumors (**Figure 4C**). No difference was observed between FRT-treated tumors and untreated control tumors at the ≥400 mm^3^ tumor regrowth end-point post-RT.

**Figure 4.**
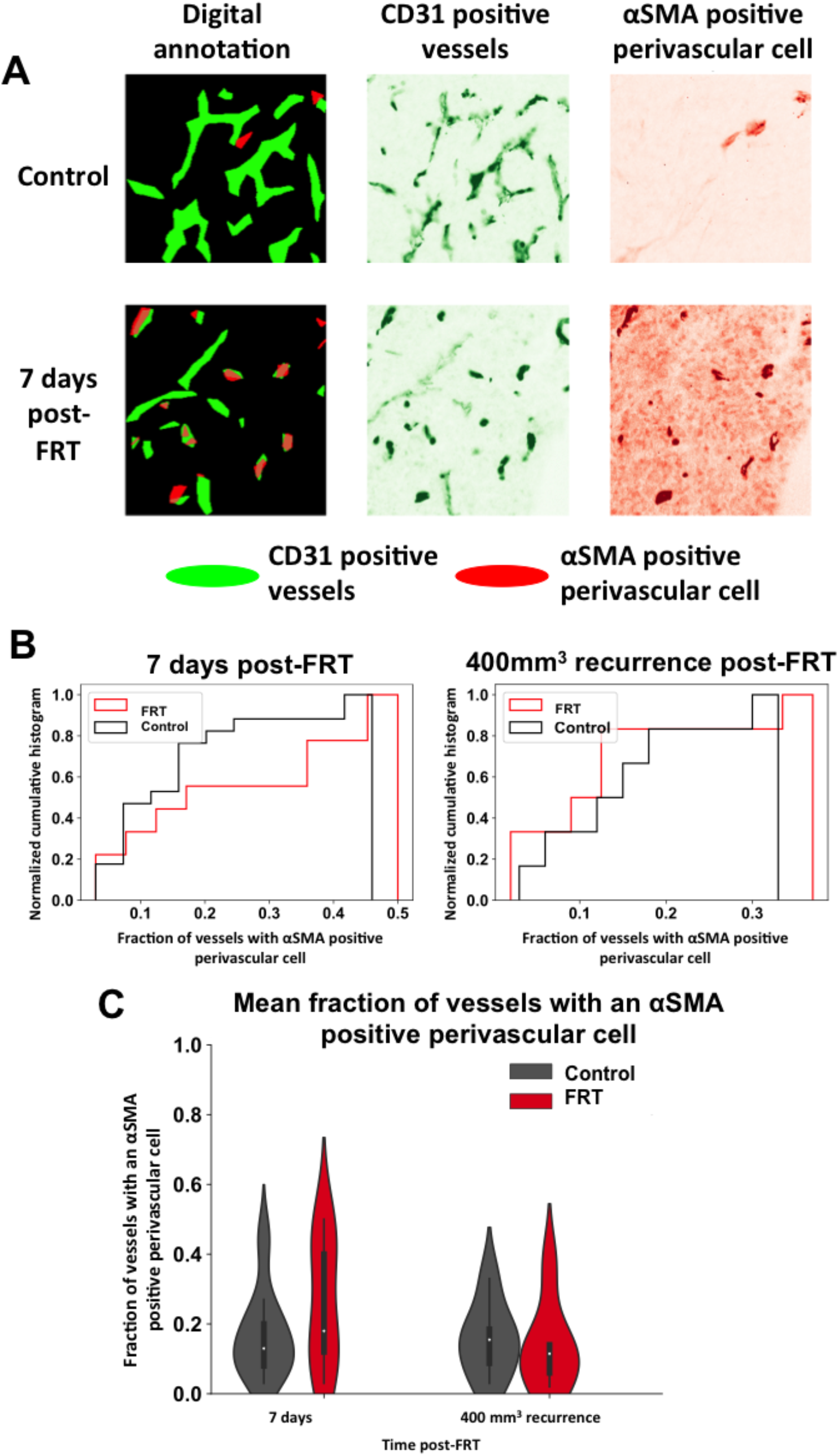
FRT induces vascular normalization in flank TRAMP-C1 PCa tumor allografts. (**A**) Image annotation analysis of immunofluorescence images from control and FRT-treated TRAMP-C1 flank tumors (green – anti-CD31; red – anti-αSMA) revealed an increased fraction of vessels with ≥1 adjacent αSMA-positive perivascular cells 7 days post-FRT (‘early’ time point). (**B**) This effect was not seen at the ≥400 mm^3^ tumor regrowth end point (‘late’ time point). (**C**) An increased mean fraction of CD31-positive tumor vessels with an adjacent αSMA-positive peri-vascular cell was observed 7 days post-FRT compared to control tumors (‘early’ time point). No difference was observed between FRT-treated and control tumors at the ≥400 mm^3^ tumor regrowth end point (‘late’ time point).

### Radiotherapy improves perfusion in prostate cancer flank tumor allografts

In order to investigate whether FRT influences perfusion in TRAMP-C1 flank tumor allografts at 7 days following initiation of FRT, dynamic contrast enhanced magnetic resonance imaging (DCE-MRI) analysis was performed (**Figure 5A**). This revealed that FRT resulted in enhanced perfusion, as measured by the iAUC at 90 seconds after Gd injection, 7 days post-FRT in TRAMP-C1 flank tumor allografts (**Figure 5B**). Gd contrast-induced enhancement of the fraction of voxels assessed by DCE-MRI was indicative of an improvement of tumor perfusion post-FRT, and this enhanced perfusion occurred predominantly in the core (the central 1/5th of the tumor segmentation by volume) of the tumor (**Figure 5B**). In order to investigate whether this FRT-induced enhanced perfusion effect could be seen using the same imaging modality in a second tumor model, this experiment was repeated using MyC-CaP tumors in FVB mice. A similar effect on tumor perfusion at 7 days post-FRT was seen in MyC-CaP tumors (**Figure 5C**). Where paired pre- and post-FRT treatment DCE-MRI iAUC90 image maps demonstrated enhanced perfusion in TRAMP-C1 flank tumor allografts, corresponding tissue samples demonstrated decreased hypoxia as demonstrated by pimonidazole staining (**Figure 5D**).

**Figure 5.**
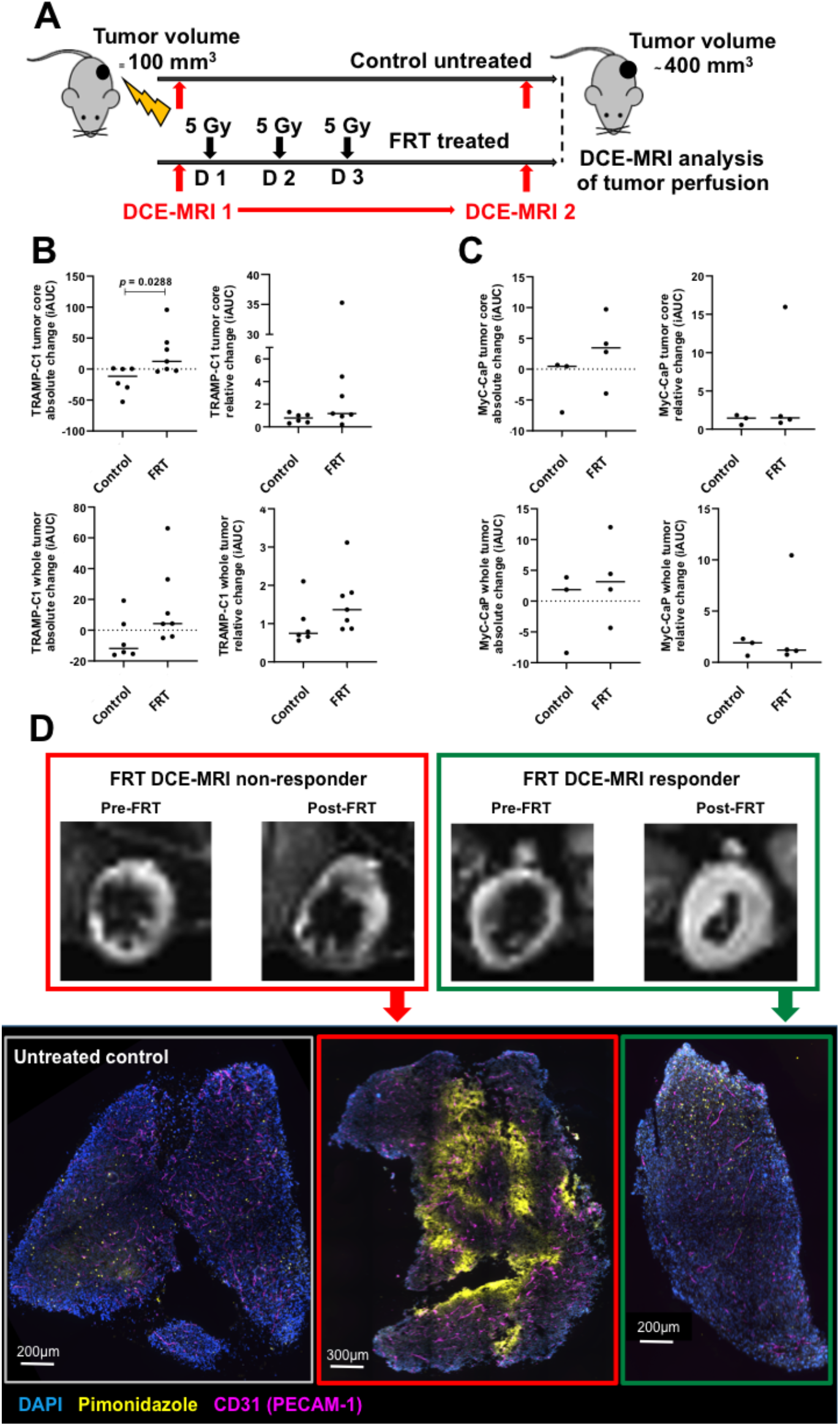
FRT improves vascular perfusion of flank TRAMP-C1 PCa tumor allografts as demonstrated using DCE-MRI. (**A**) Outline schematic of sequential DCE-MRI imaging of flank TRAMP-C1 tumor allografts immediately pre- and 7 days post-FRT. (**B**) Analysis of changes in the iAUC in paired image samples pre- and post-treatment revealed a significant increase in contrast uptake in the central core of TRAMP-C1 tumors, and a trend towards an increase in contrast uptake in the entire TRAMP-C1 tumor, following FRT versus control tumors. (**C**) Analysis of paired image samples pre- and post-FRT in MyC-CaP tumors revealed a similar trend towards an increase in contrast uptake in both the central core and the entire tumor. (**D**) Where paired pre- and post-FRT DCE-MRI iAUC90 image maps demonstrated enhanced perfusion in TRAMP-C1 flank tumor allografts, corresponding tissue samples demonstrated decreased peak hypoxia as detected by pimonidazole staining. Data are presented on a scatter plot with a line representing the median and analyzed using a two-tailed unpaired t-test (n = 6 untreated control tumors; n = 7 FRT-treated tumors). Median (range) body weight at start of experiment: untreated control tumors = 22.2 g (21.0–24.7 g); FRT-treated tumors = 21.5 g (20.0–25.6 g).

### Radiotherapy induces in vitro anti-angiogenesis of endothelial cells

The *in vivo* findings in TRAMP-C1 flank tumor allografts at 7-days post-FRT demonstrated that CD31-positive tumor vessels in this model were larger in size, fewer in number, further apart, with enhanced αSMA-positive peri-vascular cell coverage, and these tumors had enhanced vascular perfusion demonstrable on DCE-MRI. This indicated that FRT had induced vascular normalization at 7-days post-FRT in this model. In order to investigate the effects of irradiation on endothelial cell sprouting and angiogenesis *in vitro*, HUVECs were exposed to a single 2 Gy, 5 Gy or 10 Gy dose. This demonstrated that an increasing irradiation dose reduced the HUVEC cell number over time following treatment, compared against untreated control cells (**Figures 6A-C**). Using an *in vitro* hanging drop assay we observed that HUVEC endothelial cell sprouting was reduced with increasing irradiation dose at 48 hours post-treatment (**Figure 6D**).

**Figure 6.**
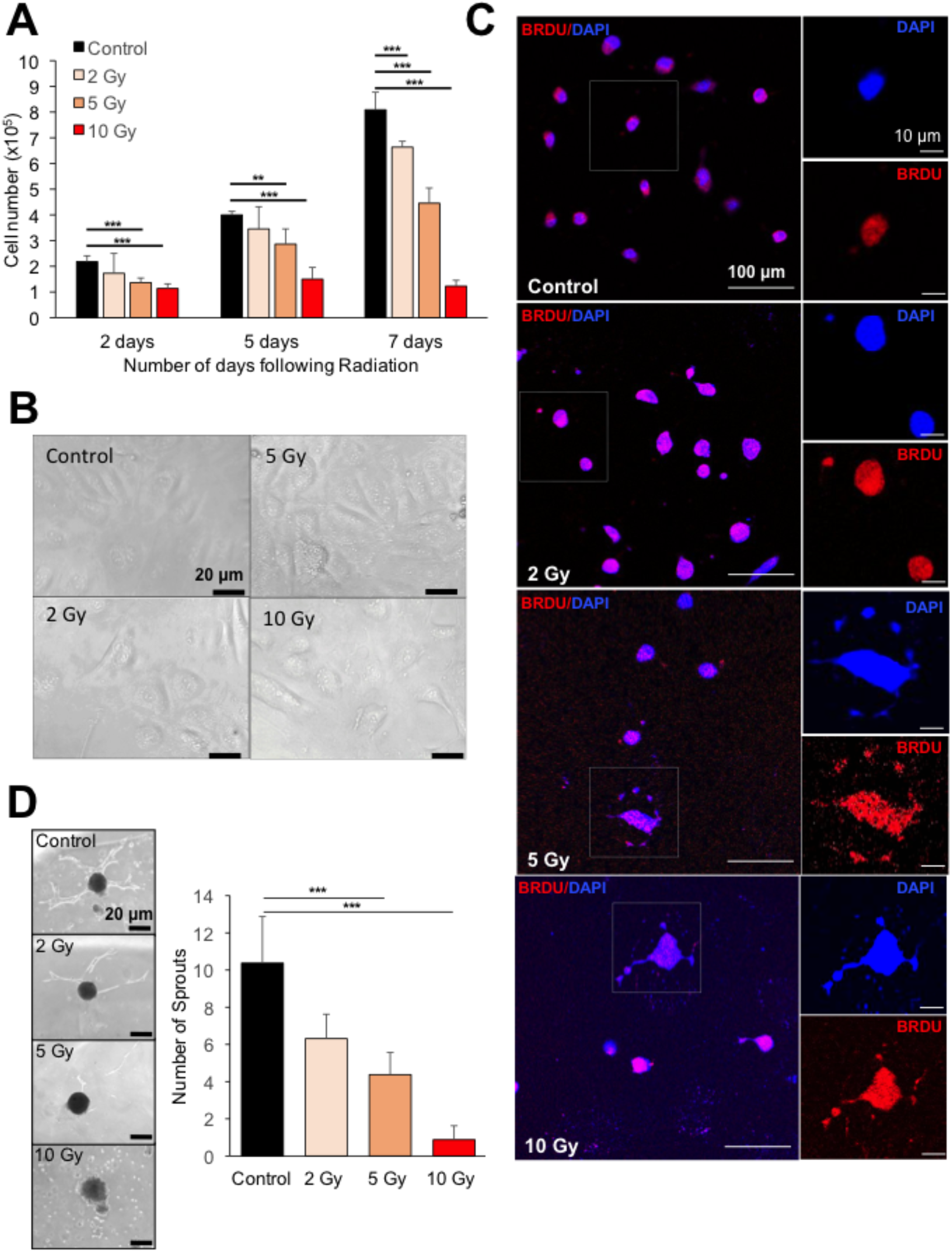
*In vitro* investigation of effects of irradiation on endothelial cell sprouting and angiogenesis. (**A-C**) A time course experiment following exposure of human umbilical vein endothelial cells (HUVECs) to 2 Gy, 5 Gy or 10 Gy irradiation demonstrated reduced HUVEC cell number, morphological cell changes, and increased number of cytoplasmic DNA fragments (DAPI, blue). (**D**) A ‘hanging drop’ assay demonstrated that endothelial cell sprouting was reduced with increasing irradiation dose at 48 hours. Error bars = standard deviation. \*\**p* <0.01, \*\*\**p*<0.0001; one-way ANOVA or unpaired Student t-test comparing two data groups; data representative of n = 3 independent experiments.

### Combination therapy of RT with VTP suppressed tumor growth to a greater extent than either treatment alone

To test the hypothesis that the vascular normalization induced by FRT might improve the outcome of VTP-mediated tumor growth control, the effects of sequential treatment of TRAMP-C1 flank allograft tumors with FRT followed by VTP were studied in syngeneic immunocompetent C57BL/6 mice (**Figure 7A**). In order to administer the VTP at the time of the observed FRT-induced vascular normalization, VTP was administered seven days following initiation of FRT (**Figure 7A**). Whilst prior studies with VTP as monotherapy had established that 7 mg/kg WST-11 had a lesser efficacy than 9 mg/kg in terms of inducing an anti-tumor effect, it was tested whether the sequential effects of FRT and VTP might enable a reduction in the WST-11 dose required for tumor growth delay. The effect of FRT followed sequentially by either 7 mg/kg or 9 mg/kg WST-11 VTP was therefore explored (**Figures 7B & 7C**). The sequential combination of FRT followed by 9 mg/kg WST-11 VTP delayed tumor growth significantly compared to 9 mg/kg WST-11 VTP alone, and led to a trend towards enhanced tumor growth delay compared to FRT alone (**Figures 7B & 7C**). The sequential combination of FRT and 7 mg/kg WST-11 VTP increased significantly tumor growth delay compared to either 7 mg/kg WST-11 VTP alone or FRT alone (**Figures 7B & 7C**). In a survival analysis, sequential combination of FRT and 9 mg/kg WST-11 VTP significantly enhanced survival to a tumor volume of 400 mm^3^ compared to 9 mg/kg WST-11 VTP alone or FRT alone (**Figure 7D**). Similarly, the sequential combination of FRT and 7 mg/kg WST-11 VTP significantly enhanced survival to a tumor volume of 400 mm^3^ compared to 7 mg/kg WST-11 VTP alone or FRT alone (**Figure 7C**). Sequential combination of FRT and 7 mg/kg WST-11 led to the longest survival to 400 mm^3^ and a long-term cure in one animal (**Figure 7D**), suggesting that the lower 7 mg/kg dose of WST-11 may have greater efficacy when combined with FRT than the 9 mg/kg WST-11 dose.

**Figure 7.**
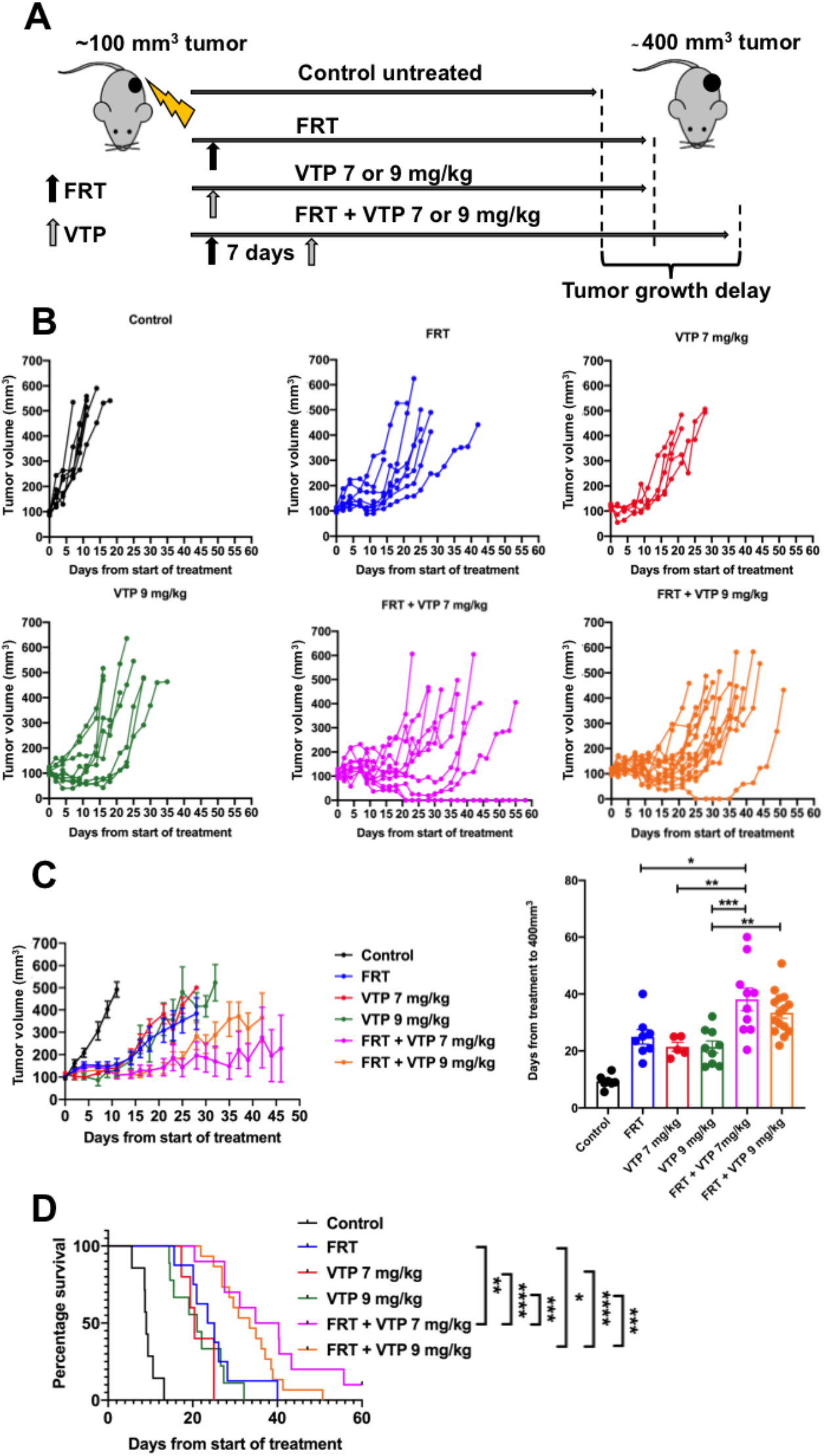
Multimodality therapy with FRT followed at 7 days by VTP causes tumor growth delay in TRAMP-C1 flank tumor allografts. (**A**) Outline schematic of treatment of tumors with FRT, VTP (7 or 9 mg/kg WST-11), or a sequential combination of FRT followed at 7 days by VTP (7 or 9 mg/kg WST-11). (**B**) Tumor growth delay analysis of tumors following treatment with FRT, VTP (7 or 9 mg/kg WST-11), or sequential combined FRT and VTP (7 or 9 mg/kg WST-11). (**C**) Sequential combined FRT and VTP (7 or 9 mg/kg WST-11) significantly delayed tumor growth compared to either FRT or VTP alone. (**D**) Mice treated with sequential FRT and VTP (7 or 9 mg/kg WST-11) had significantly improved survival to tumor regrowth end-point of 400 mm^3^ compared to treatment with either FRT or VTP alone. Numbers per group: control n = 7, FRT n = 8, VTP 7 mg/kg n = 5, VTP 9 mg/kg n = 9, FRT and VTP 7 mg/kg n = 10, FRT and VTP 9 mg/kg n = 15. Median (range) body weight at treatment: control = 21.1 g (20.8 – 22.1 g), FRT = 21.1 g (19.8 – 24.3 g), VTP 7 mg/kg = 21.6 g (20.8 – 23.8 g), VTP 9 mg/kg = 20.9 g (18.5 – 23.8 g), FRT and VTP 7 mg/kg = 21.9 g (18.9 – 24.3 g), FRT and VTP 9 mg/kg = 21.3 g (19.6 – 23.8 g). Data in treatment groups are presented as individual tumor growth kinetics (**B**), grouped tumor growth kinetics (**C**), mean ± SEM growth delay to ≥400 mm^3^ (**C**), and survival to ≥400 mm^3^ using Kaplan-Meier curves (**D**). Data were analyzed using ordinary oneway ANOVA with Tukey’s*post hoc* adjustment for multiple comparisons (**C**), and Log-Rank (Mantel-Cox) test (**D**). * *p*<0.05; ** *p*<0.01; *** *p*<0.001; **** *p*<0.0001.

Finally, the hypothesis that tumor growth delay for treated TRAMP-C1 flank allograft tumors was enhanced for VTP when sequentially delivered following initial FRT, compared to VTP alone, was tested. When analyzing the time for tumor regrowth from 150mm^3^ at VTP delivery to the 400 mm^3^ end-point, a trend was observed towards enhanced delay in tumor growth for VTP following previous FRT, compared to VTP alone for the 9 mg/kg WST-11 (**Figure 8A**) and 7 mg/kg WST-11 (**Figure 8B**) doses. A combined analysis of mice treated with 7-9 mg/kg WST-11 VTP alone versus combined sequential FRT and 7-9 mg/kg WST-11 (**Figure 8C**) showed a statistically significant enhancement in tumor growth delay following administration of VTP if it was delivered 7 days following FRT, compared with VTP alone.

**Figure 8.**
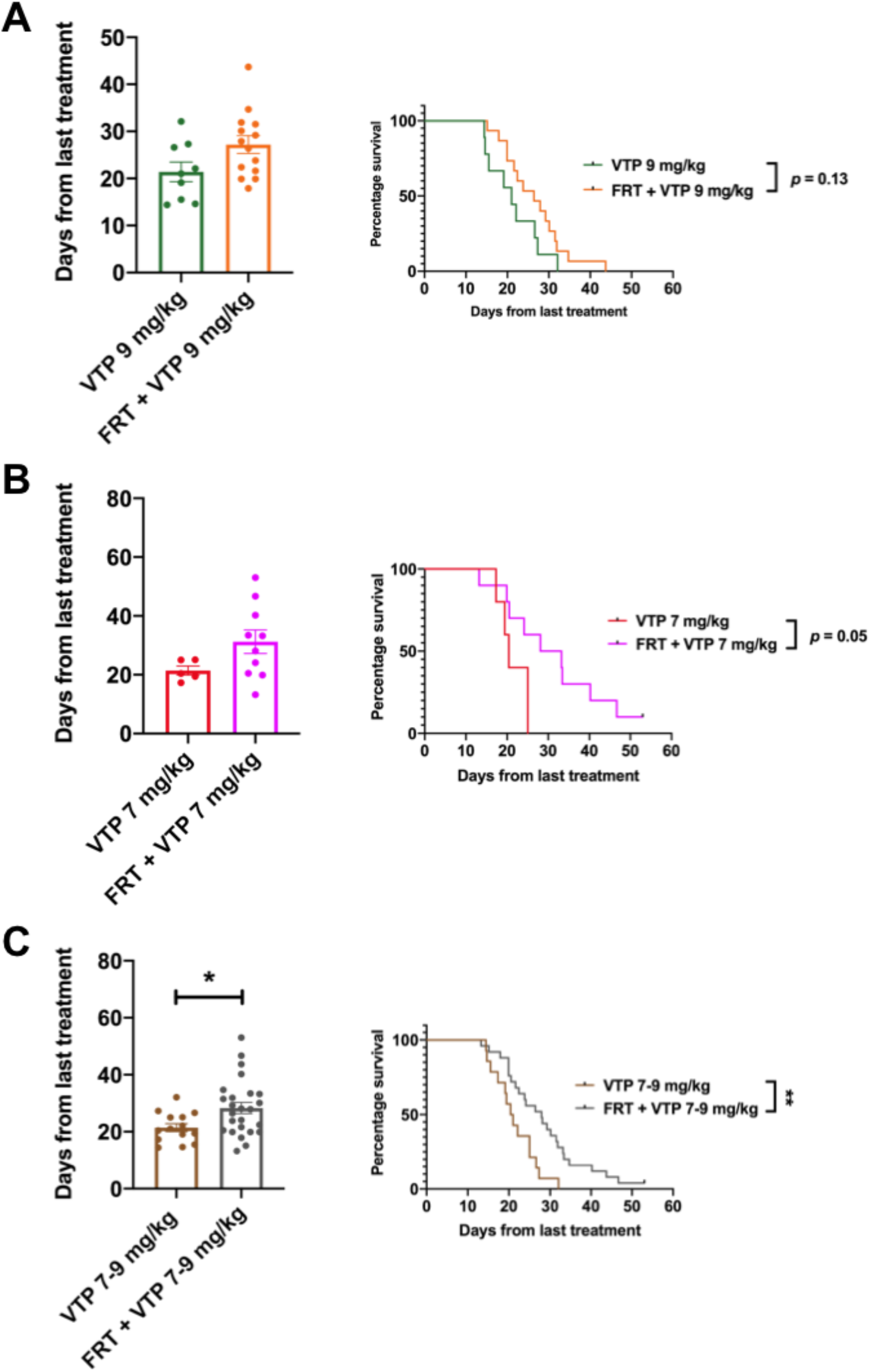
Neo-adjuvant FRT improves efficacy of subsequent VTP delivered 7 days later. Analysis of post-VTP tumor growth delay of TRAMP-C1 flank allograft tumors treated with VTP 7 days post-FRT, compared to VTP alone. (**A-B**) Analysis of time for tumor regrowth from 150 mm^3^ at VTP delivery to the 400 mm^3^ end-point revealed a trend towards enhanced tumor growth delay for VTP post-FRT compared to VTP alone for 9 mg/kg WST-11 and 7 mg/kg WST-11. (**C**) Combined analysis of mice treated with 7-9 mg/kg WST-11 VTP alone versus combined sequential FRT and 7-9 mg/kg WST-11 revealed a significantly enhanced tumor growth delay following administration of VTP if delivered 7 days post-FRT, compared with VTP alone. Data were analyzed using a two-tailed unpaired t-test for the bar chart and Log-Rank (Mantel-Cox) test for the Kaplan-Meier curves. * *p*<0.05; ** *p*<0.01.

## Discussion

There is an unmet clinical need to increase the cure rate of high risk PCa, and reduce treatment toxicity. Patients with these grades/stages of PCa are typically offered either radical surgery or radical FRT (the latter with neoadjuvant and concomitant ADT for up to three years). However, around a third of patients develop disease recurrence, and treatments lead to significant side effects on urinary, sexual and bowel function, as well as loss of quality of life in long-term survivors. There is an increasing interest in the role of multimodality therapy for PCa (37). To date PCa is one of few malignancies where immunotherapy is not part of standard of care (38), potentially due to the immunosuppressive tumor microenvironment (39) and/or its low mutational burden (40,41). It is likely that other treatment combinations beyond immunotherapy are necessary to improve PCa tumor control, and reduce treatment toxicity.

VTP is a minimally invasive focal ablation precision surgical technique, largely evaluated as monotherapy for low-volume low- or intermediate-risk PCa (26–28,42–44). The technique has not been evaluated clinically as part of a multimodality therapy approach, or in high-risk PCa. Importantly the available evidence from several clinical trials demonstrated that VTP has a good safety and tolerability profile (26–28), with a return to baseline urinary and sexual function by 6 months post treatment. Given that the treatment parameters for VTP in terms of WST-11 photosensitizer dose and activation have been established for optimal targeted tumor ablation, along with the fact that FRT for PCa is well established, it could be feasible, based on pre-clinical data, to combine these treatments in early and subsequent late phase clinical trials. Herein, we provide evidence that initial moderate hypofractionated FRT ahead of VTP, with VTP being delivered during a window of time when the irradiated PCa tumor displays vascular normalization, leads to enhanced delay in tumor growth, with a resultant survival benefit in mice and the possibility of complete tumor cure. It may be possible, and indeed it is attractive, to combine FRT with VTP in treating PCa. This may obviate the need for neoadjuvant and/or concomitant ADT, as well as reduce the dose of conventional FRT administered. This new multimodality therapy approach will require formal clinical evaluation. The conception of an organ-sparing strategy in PCa management, which is directed towards a focal lesion or takes a hemigland approach in order to deliver effective tumor control and/or cure with reduced toxicity compared to whole gland treatment, is becoming an increasing focus of research for PCa as it may lead to an improved quality of life compared against current standard of care whole gland therapy (42,45,46). VTP has distinct advantages as a form of focal therapy; the technique has been evaluated in phase-III randomized clinical trial (28), and it can anatomize treatment such that it conforms to the malignant lesion along with a margin of normal tissue within the prostate gland, preserving urinary and sexual function. Pre-clinical studies in PCa models have demonstrated that VTP may be successfully combined with ADT (47), a ^111^In-DOTA-AR bombesin antagonist (48), and anti-macrophage colonystimulating factor (anti-CSFR1) (49), however VTP has not been investigated in close sequential combination with FRT. VTP has been demonstrated to be safe and effective in patients with recurrent PCa following FRT however the patients in these early phase clinical trials received FRT and VTP with a considerable interval of several years. The evidence we present here suggests that sequential combined FRT and VTP within a short interval enhances anti-tumor control and warrants evaluation in clinical trials.

The observation that VTP delivered shortly after initial moderate hypofractionated RT increases delay in tumor growth is intriguing, and suggests that neo-adjuvant FRT delivered in the correct fraction size and dose improves the efficacy of VTP. One possible explanation is that the transient vascular normalization induced by moderate hypofractionated RT, with the resultant transient increase in tumor vessel perfusion, leads to enhanced WST-11 delivery into the tumor, such that targeted near-infrared illumination of the target tumor lesion is then more effective in terms of tumor ablation. We observed that FRT enhances tumor perfusion as observed on DCE-MRI 7 days post-FRT, and this is consistent with other studies using similar fractions and doses of FRT (50). The dose and fraction size of FRT, along with the window of time in which to look for vascular normalization, are key issues as it is known that large irradiation doses rapidly destroy the tumor vasculature, rather than lead to transient enhanced perfusion and function (34,51–55). The results of *in vivo* xenograft pre-clinical model experiments show that the size and number of irradiation fractions determines whether transient induction of enhanced tumor vascularity occurs. This can be generalized as follows: 5-10 Gy fractions cause transient blood flow increase, which returns to normal; 10-15 Gy fractions cause tumor blood flow decrease, which then recovers; 15-20 Gy fractions cause blood flow to rapidly decrease, and vessels deteriorate (53,56). It is important for future clinical translation to consider these aspects of FRT delivery. Hypofractionated RT is increasingly used clinically (37), and it would be entirely feasible to use FRT schedules that are likely to induce vascular normalization ahead of VTP in a multimodality approach.

We observed that the lower 7 mg/kg dose of WST-11 resulted in a more pronounced reduction in tumor growth following FRT than the 9 mg/kg WST-11 dose. It is possible that near-infrared illumination of 7 mg/kg WST-11 ablates fewer tumor vessels than 9 mg/kg WST-11, leaving sufficient tumor vessels available for immune cells to infiltrate the tumor and elicit an anti-tumor immune response, thereby enhancing the observed anti-tumor effects.

The concept of targeting the tumor vasculature, rather than the cancer cells, has been explored in a large number of pre-clinical and clinical studies. Tumor vessels are often functionally abnormal (31,32) potentially rendering them susceptible to targeted therapy with molecular agents. In the case of VTP, which requires a functional vasculature, it is possible that prior enhanced vascular function through vascular normalization (33) improves anti-tumor effects of subsequent VTP treatment. A variety of vascular targeting agents have been evaluated in pre-clinical cancer research, including inhibitors of angiogenesis and vascular disrupting agents (VDAs). Some VDAs have been investigated in combination with FRT (57–62), where the VDA was administered following initial FRT. Combining VDAs with conventional therapies such as FRT may improve treatment outcomes by enhancing antitumor efficacy, with non-overlapping toxicities, and spatial cooperation. VTP may have particular clinical benefit as a VDA compared to drugs such as DMXAA, CA4DP and ZD6126 investigated in other studies. WST-11 used for VTP has minimal toxicity as it is focally activated by near-infrared illumination in the tumor vasculature rather than being active systemically. VTP is therefore a more attractive clinical agent than other VDAs in combination with FRT.

This study has several limitations. Firstly, it has not investigated the immune response to sequential FRT and VTP. FRT (36) and VTP (24,25) each induce immunological changes within tumors including PCa. VTP may induce immunogenic cell death and thereby auto-vaccinate a patient against his PCa lesion. The effects of sequential FRT and VTP on the tumor immune microenvironment are being investigated separately. Secondly, our work was performed on the TRAMP-C1 flank tumor allograft model of PCa in C57BL/6 mice. This is the most commonly used *in vivo* pre-clinical PCa model, and it would be helpful to validate the benefit of combined sequential FRT and VTP in an alternative model of PCa, although the feasibility of combining FRT and VTP in an orthotopic small animal model such as mouse would be challenging in view of concerns over rectal and urethral toxicity of the combined treatment. Thirdly, the experiments described have not incorporated ADT, which is conventionally used in the neo-adjuvant clinical setting prior to FRT. ADT could modulate the effect of sequentially combined FRT and VTP. Fourthly, although we demonstrate that the growth kinetics of TRAMP-C1 allograft tumors are slower post-VTP following prior FRT, compared to VTP monotherapy, it will be important to investigate the microenvironment of recurrent tumors, as recurrence following combination therapy may result in more aggressive disease. Fifthly, we only administered a single VTP treatment following FRT, whereas repeat VTP can be administered. Finally, given that a functional tumor vasculature and viable endothelial cells after FRT are conventionally considered to be undesirable post-treatment effects as they can promote tumor recurrence (63) and metastasis (64), the clinical safety of delivering VTP during a window of vascular normalization following sublethal FRT needs to be assessed further.

In summary, these findings demonstrate that combining FRT and VTP may be a promising clinical strategy in the treatment of PCa. These results could pave the way for the development of future early phase clinical trials in patients with high-risk localized and/or locally advanced disease.

## Materials and Methods

### Cell lines and cell culture

TRAMP-C1 (ATCC^®^ CRL-2730™) and MyC-CaP (ATCC^®^ CRL-3255™) cells were purchased from American Type Culture Collection (ATCC^®^) and cultured as previously described (36). Human umbilical vein endothelial cells (HUVECs) (up to passage 6; Lonza, Wokingham, UK) were cultured in EGM-2 medium (Lonza, Wokingham, UK).

### Endothelial growth changes

HUVECs were irradiated with a ^137^Caesium irradiator (IBL 637, CIS Bio International). Cells were exposed to single 2 Gy, 5 Gy or 10 Gy dose of irradiation. Viable cells (excluding trypan blue positive dead cells) were counted over a time course of seven days to assess changes in HUVEC growth. Images of cells *in situ*, using a white-light microscope, were taken 48 hours after RT.

### Endothelial DNA damage assessment

Following 2 Gy, 5 Gy or 10 Gy irradiation with a ^137^Caesium irradiator HUVECs were seeded onto glass slides. 48 hours post-RT cells were stained with 4,6-diamidino-2-phenylindole (DAPI; Invitrogen). The slides were washed twice with PBS and mounted with Vectashield (Vector Labs). DNA damage was assessed using confocal microscopy (Zeiss 780 inverted confocal microscope) and 405 nm fluorescence excitation.

### Endothelial sprouting assay

Following irradiation of HUVECs with single 2 Gy, 5 Gy or 10 Gy doses using a ^137^Caesium irradiator endothelial cell spheroids were generated by the hanging drop method as previously described (65).

### In vivo study approval

Animal procedures were performed according to UK Animal law (Scientific Procedures Act 1986) and ARRIVE guidelines, with local ethics and Home Office approval.

### Generation of flank tumor allograft model

Naive 6–8 week old male immunocompetent C57BL/6 and FVB mice (Charles River Laboratories, UK) were housed in groups of six in a pathogen-free facility with 12 hour light cycles, in individually ventilated cages on woodchip bedding, with access to water and food *ad libitum*, at 22°C (range 21– 24°C) and 50% humidity (range 35–75%), with environmental enrichment and bedding material, and monitored for body weight changes twice weekly (36). 2 x 10^6^ TRAMP-C1 (C57BL/6 mice) or 1 x 10^6^ MyC-CaP (FVB mice) cells in PBS and 1: 1 high concentration phenol red-free Matrigel^®^ (Corning) were injected into the flank of mice under isofluorane inhalational anesthesia on a heat mat. Tumors were measured in the home cage pre- and post-treatment using digital callipers three times per week (tumor volume = π/6 x length x width x height). When tumors reached 100-120 mm^3^ mice were assigned to treatment groups on a first come, first allocated basis using a randomly generated treatment list (GraphPad Prism 8, GraphPad Software, USA). To detect tumor hypoxia, mice were injected intraperitoneally with 60 mg/kg pimonidazole solution 90 minutes before euthanasia. All mice were culled as described previously (36).

### Radiotherapy

FRT was delivered as previously described (36).

### Dynamic contrast enhanced magnetic resonance imaging

Inhalational anesthesia was induced and maintained with isofluorane (1–4% in air) in order to maintain a respiration rate of 40-60 breaths per minute, and temperature was maintained at 35°C using a homeothermic temperature maintenance system as previously described (50,66,67). Dynamic contrast enhanced MRI (DCE-MRI) was performed at 4.7 and 7.0T scanners (Varian, VNMRS console) using 25 mm inner diameter birdcage coils (Rapid Biomedical, Germany) as previously described (50). Respiratory-gated 3D gradient echo scans (echo time = 0.6 ms; repetition time = 1.15 ms; nominal 5 degree flip angle) with an isotropic resolution of ~420 μm and a respiratory rate dependent frame acquisition time of ~8-10 seconds were obtained (50). Fifty frames were acquired with a 25 μl bolus infusion of Gadolinium (Gd) solution (Omniscan GE HEALTHCARE) administered using a syringe pump (PHD2000, Harvard Apparatus) over 5 seconds commencing at the beginning of frame 11. Radiofrequency field inhomogeneities were accounted for using a respiratory-gated implementation of the Actual Flip Angle technique, and baseline T1 was measured using a variable flip angle sequence as reported previously (50). In order to analyze tumor perfusion a manual segmentation was initially performed from the average image of the DCE sequence using ITK-SNAP medical image segmentation software (68). The MR signal was then converted to Gd concentration as previously described (69). The initial area under the Gd curve (iAUC) was measured by integrating the first 90 seconds after injection, and used as an indicator of perfusion.

### VTP administration

In order to administer VTP treatment in our small animal facility, we constructed a bespoke optical excitation system. A small enclosure (**Figure 1**) contained an animal heating pad, tubing for inhalational anesthesia, and physiological monitoring apparatus. Animal imaging, illumination and fibre-coupled excitation and guide beam illumination optics were housed in the lid of this enclosure. WST-11 VTP excitation was provided by a 755 nm thermoelectrically-cooled semiconductor laser diode (LDX Optronix, Missouri, US, type USA LDX-3210-750), and its output power was controlled by bespoke hardware controlled by a software system and a graphical user interface, which also controlled optical exposure time. A low power (<1 mW) excitation guide beam was provided, generated by a single 590 nm LED (type SMB1N-590-02, Roithner LaserTechnik GmbH, Austria), the output of which was combined with the laser output using a dichromatic reflector (DMLP638R, Thorlabs, UK) prior to launching into the output fibre. A multimode fibre (type M53L02 Ø600 μm, 0.50 NA, Thorlabs, UK) carried both the guide beam and excitation light to the enclosure where it was homogenized and collimated to a slightly diverging beam diameter of 8 mm nominal, delivering a typical excitation intensity of 120 mW/cm^2^. A simple mechanical system and turning prism allowed the excitation and guide beam to be positioned over the area of interest. Additionally the enclosure was fitted with diffuse LEDs operating at 590 nm (Roithner LaserTechnik GmbH, Austria type SMB1N-590-02). The animal in the enclosure could be viewed at all times with the aid of a miniature high dynamic range monochrome camera mounted in the enclosure lid. The wavelength of the guide beam and the illumination (590 nm) was chosen as it was in the nadir of the WST-11 VTP spectral absorption. Similarly, WST-11 injections were performed under yellow light generated by an array of T1¾ 590 nm LEDs (type HLMP-EL3G-VX0DD, Broadcom, USA). Lyophilized WST-11 was a kind gift from Professor Avigdor Scherz, Weizmann Institute of Science, Israel, and was reconstituted in sterile 5% dextran in water at 2 mg/mL under light protected conditions, and aliquots were stored at −20°C in the dark. Aliquots were thawed on the day of VTP treatment and sterile filtered through a 0.2 μm disc syringe filter. Mice with tumors measuring 120 mm^3^ received VTP treatment in the morning under isofluorane inhalational anesthesia with physiological monitoring using a bespoke cradle. Anaesthetized mice received an intravenous infusion of 7 or 9 mg/kg WST-11 via the tail vein followed immediately by 10 minutes laser excitation of the subcutaneous flank tumor at an intensity of 120 mW/cm^2^ using a collimating lens. The optical irradiation light field was arranged to cover the entire subcutaneous flank tumor area plus a 1 mm rim. Mice were returned to the home cage following recovery from anesthesia and underwent post-procedure health monitoring along with tumor measurements using digital callipers.

### Immunofluorescence of tissue sections

Freshly excised TRAMP-C1 tumors were placed into a cryomold filled with optimal cutting temperature compound and quickly frozen in a cold bath containing isopropanol and dry ice. 10 μm frozen tissue sections were cut and placed onto super frost microscope slides (FisherBrand), and fixed with 4% dilution electron microscopy grade paraformaldehyde (Electron Microscopy Sciences) in PBS for 30 minutes at room temperature in a Coplin jar. Tissue section slides were then washed with PBS in a Coplin jar prior to a combined blocking and permeabilization step using 3% BSA in 0.3% triton/PBS for 90 minutes at room temperature. Tissue sections were then encircled with a hydrophobic PAP marking pen, and 200 μl of primary antibody (1:200 anti-αSMA antibody, eBioscience, clone 1A4, efluor 570; 1:200 anti-CD31 antibody, Biolegend, clone MEC13.3, Alexa Fluor 488; 1:100 anti-pimonidazole antibody, Hypoxyprobe plus Kit, clone 4.3.11.3, FITC) diluted in blocking solution was added for incubation in a humidified chamber at 4°C overnight, with this and all subsequent steps under light protection. After removal of primary antibody a 300 nM DAPI nuclear counterstain in PBS was applied for 30 minutes incubation prior to washes in PBS, and mounting with coverslips sealed with Wirosil dental silicone. αSMA and CD31 sample images were acquired with a Plan Apo 20x 0.75NA objective on a Nikon Ti Widefield Microscope equipped with a Lumencor SpectraX LED light source and an Andor Neo Zyla Camera. Pimonidazole sample images were acquired using a Leica DM18 inverted widefield microscope and a Hamamatsu Orca Flash 4.0 V3 camera, a Lumencor SOLA-SE-II Light source (365nm) and HC PL APO 20x/0.80 lens.

### Digital analysis of tumor vessels

Tumor CD31-positive blood vessel quantification was performed with classical image processing methods. Following image pre-processing, image segmentation was performed to identify tissue areas and CD31-positive blood vessels in an automated fashion. The vessel segmentation was analysed using the Euclidian distance transformation to obtain the local thickness of all segmented objects in the images. The combined binary masks indicating the presence of tissue and CD31-positive blood vessel masks were used to compute the vessel density and the distance from any tissue pixel to the closest vessel.

### Digital annotation and quantification of tumor vessels and αSMA-positive peri-vascular cells

Computer-assisted image annotation was used to identify αSMA-positive perivascular cells. Since αSMA is not exclusively expressed in pericytes, αSMA-positive perivascular cells were annotated by overlapping the image channels corresponding to αSMA (red) and CD31 (green), and only selecting αSMA-positive peri-vascular cells adjacent to, or overlapping, CD31-positive blood vessels. To assist annotation, αSMA-positive cells that had a distance of more than 10 pixels, or 3.2 μm, were suppressed. Annotation was performed using the Annotation of Image Data by Assignments (AIDA) web application platform (70) developed in-house for the annotation of large microscopy images. Here, relevant regions of interest in the tumor were selected, and the annotators then marked αSMA-positive peri-vascular cells and CD31-positive blood vessels in these regions. Areas in the αSMA channel selected by at least 3 of 4 independent observers were accepted as αSMA-positive peri-vascular cells. In order to investigate the extent of αSMA-positive perivascular cell coverage of CD31-positive blood vessels, the fraction of blood vessels in each region presenting spatially adjacent αSMA-positive perivascular cells was computed. Control untreated samples were compared against FRT-treated samples.

### Statistical analysis

Statistical analysis was performed using GraphPad Prism 8 (GraphPad Software, USA). For *in vitro* work ordinary one-way ANOVA tests were performed with Dunnett’s or Tukey’s post hoc adjustment for multiple comparisons. For *in vivo* tumor growth delay experiments, ordinary one-way ANOVA was performed using Tukey’s test for multiple comparisons following Brown-Forsythe’s test for equality of the means. Tumor growth delay was defined as a significant increase in time (days) for a tumor treated at 100-120 mm^3^ to reach end-point size of 400 mm^3^ compared against control untreated tumors, a single mouse being considered an experimental unit. All results are mean ± standard error of the mean. *P*<0.05 was considered to be a statistically significant difference.

## Author Contributions

Conception and design: RJB, RJM, AS, FCH.

Development of methodology: HTS, YP, AM, IDT, EB, AC, JL, KHT, JR, SCS, PDA, IGM, AH, BV, RJM, AS, FCH, RJB.

Acquisition of data (provided animals, acquired and managed patients, provided facilities, etc.): HTS, YP, AM, EB, AC, JL, KHT, EM, SCS, PK, SG, PDA, DAS, RJB.

Analysis and interpretation of data (e.g., statistical analysis, biostatistics, computational analysis): HTS, YP, EB, AC, JL, KHT, PDA, RJB.

Writing, review, and/or revision of the manuscript: HTS, YP, AM, IDT, EB, AC, JL, KHT, EM, JR, DP, LA, TY, SCS, PK, SG, PDA, DAS, SJL, DAW, ADL, IGM, AH, RJM, BV, AS, FCH, RJB. Administrative, technical, or material support (i.e., reporting or organizing data, constructing databases): HTS, YP, AM, IDT, EB, AC, JL, KHT, EM, JR, DP, LA, TY, SCS, PK, SG, PDA, DAS, SJL, DAW, AH.

Study supervision: AH, RJM, BV, AS, FCH, RJB.

## Acknowledgements

This work was funded by a Cancer Research UK/Royal College of Surgeons of England Clinician Scientist Fellowship (reference C39297/A22748, for RJB and HS), and by a research grant from The Urology Foundation (for YP), along with a Cancer Research UK Clinical Research Training Fellowship (for YP). BV and IDCT were funded by a Cancer Research UK program grant (C5255/A12678). JL was supported by a FRQNT postdoctoral scholarship (reference 257844). AM was funded by the John Fell Fund and the John Black Charitable Foundation. The authors also acknowledge Federal funds from the National Cancer Institute, National Institutes of Health, under Contract No. 75N91019D00024. The content of this publication does not necessarily reflect the views or policies of the Department of Health and Human Services, nor does mention of trade names, commercial products, or organizations imply endorsement by the U.S. Government.

